# Analytical solutions for the short-term plasticity

**DOI:** 10.1101/2023.04.03.535315

**Authors:** Paulo R. Protachevicz, Antonio M. Batista, Iberê L. Caldas, Murilo S. Baptista

## Abstract

Synaptic dynamics plays a key role in neuronal communication. Due to its high-dimensionality, the main fundamental mechanisms triggering different synaptic dynamics and its relation with the neurotransmitters release regimes (facilitation, biphasic, and depression) are still elusive. For a general set of parameters, and by means of an approximated solution for a set of differential equations associated with a synaptic model, we obtain a discrete map that provides analytical solutions that shed light into the dynamics of synapses. Assuming that the presynaptic neuron perturbing the neuron whose synapse is being modelled is spiking periodically, we derive the stable equilibria and the maximal values for the release regimes as a function of the percentage of neurotransmitter released and the mean frequency of the presynaptic spiking neuron. Assuming that the presynaptic neuron is spiking stochastically following a Poisson distribution, we demonstrate that the equations for the time average of the trajectory are the same as the map under the periodic presynaptic stimulus, admitting the same equilibrium points. Thus, the synapses under stochastic presynaptic spikes, emulating the spiking behaviour produced by a complex neural network, wander around the equilibrium points of the synapses under periodic stimulus, which can be fully analytically calculated.

**Author summary:** Based on the model proposed by Tsodyks et al., we obtained a map approximation to study analytically the dynamics of short-term synaptic plasticity. We identified the synaptic regimes named facilitation, depression, and biphasic in the parameters space, and determined the maximal and equilibrium points of active neurotransmitters for presynaptic neurons spiking periodically and stochastically following a Poisson process. Besides that, we verify that the time average of the variables for the synaptic dynamics driven by presynaptic neurons spiking following a Poisson distribution presents the equilibrium points obtained for the synaptic driven by periodic presynaptic neurons, spiking with a frequency that is the mean frequency of the Poisson distribution. These results shed analytical light into the understanding of synaptic dynamics.

## Introduction

Synapses are specialized structures in neuronal communication which play a key role in the transmission of neuronal signals in the brain [1]. There are two types of synapses, the electrical and the chemical [2]. Through the electrical synapses, neurons communicate to each other by direct exchange of ion currents [3]. In the axon terminals of the presynaptic neurons with chemical synapses, the action potentials generate the release of neurotransmitters in the synaptic cleft that arrive in the receptors and then produce a current in the postsynaptic neurons [4]. In the mammalian brain, most synapses are chemical [5]. The effectiveness of these transmitted currents depends on the synaptic strength that usually changes in time due to the previous activity of the synapse [6].

Some mathematical models were proposed in the literature to describe the dynamics of chemical synapse. Many of them provide a simplified description about the signal transmission between the neurons [7], for instance, single exponential decay, alpha, and double exponential synaptic functions [8, 9]. However, in these models, the maximal conductance associated with each spike event has a fixed value and does not exhibit synaptic regimes, such as facilitation and depression [10]. Others, however, provide more realistic synaptic dynamics where the intensity of synaptic conductance is altered over time [11]. This process of change in the synaptic intensity is called synaptic plasticity [12], where short- and long-term are the two main types of synaptic plasticity [13]. While for a long time scale, plasticity can exhibit long-term potentiation (LTP) or depression (LTD) [14, 15], for short time scales, the synaptic dynamics are associated with facilitation, depression and biphasic regimes [16–18]. Both facilitation and depression synaptic regimes are found between excitatory neurons in the neocortex [19]. These two kinds of plasticity also have different functions in the brain, for example, long-term plasticity is associated with processes such as memory and learning, while short-term plasticity is related to processing functions and working memory in neural circuits [20, 21].

In simple terms, short-term plasticity consists in the changes of synaptic strength conductance in a relatively small period of time which are associated with the release of neurotransmitters in each synapse [22]. These synaptic changes can act in the time scale of milliseconds to seconds, but can also last longer in the order of minutes [23–25]. A relevant model in this framework for short-term plasticity was introduced by Markram et al. [26]. According to it, spiking frequency and the amount of neurotransmitters released in the synaptic cleft are two important factors in synaptic dynamics [22]. The mechanism described by the mentioned model considers that the changes in synaptic transmission strengths depend mainly on the spike activity of the presynaptic neuron [27]. Depending on the time constants, frequency of spikes, and the amount of neurotransmitter released, different regimes as facilitation, depression, and biphasic regime of the synapse can emerge [28]. The synapses with a high probability of release tend to present short-term depression [29]. On the other hand, facilitation regimes emerge when the number of vesicles available increases due to consecutive spikes [30]. In addition, the combination of depression and facilitation has been reported to generate particular synaptic and network dynamics [31, 32].

An important behaviour found in neuron activities is the spike variability [7]. To study neuronal activities, Poisson processes are a standard to model the spike time variability of irregular firings [33–35]. The Poisson process can be considered as homogeneous [36] or inhomogeneous [37]. The main difference between these two types of Poisson processes is that the homogeneous has a constant rate of events over time while the inhomogeneous has the rate of events changing over time [38, 39]. In the field of neuroscience, the Poisson process is considered as an approximation for spontaneous neuron spikes that can be useful to investigate some aspects of neuronal dynamics [40–42]. Neurons can exhibit near periodic and Possonian spike activities over time [43]. The comprehension of how periodic and Poissonian spike activities induce different synaptic dynamics can shed light into the understanding of neuronal communication.

In this work, we obtain a map from a synaptic model described by a set of fourth ODEs proposed by Tsodyks et al. [22], and analyse how different synaptic states depend on relevant parameters, such as the spiking frequency of presynaptic neuron and the percentage of neurotransmitters released. We obtain analytically the equilibrium points in the periodic regime, identify the synaptic regimes, and determine the final and maximal value of active neurotransmitters in the space parameter of frequency and the fraction of neurotransmitters released. The maximal value corresponds to the most intense release of neurotransmitters that can generate the highest synaptic currents on the postsynaptic neurons. However, the interest in analytically calculating these values is because the solution will depend on the peculiarities of the transient behaviour in biphasic and depression, and the asymptotic behaviour of the facilitation. Furthermore, we identify when the maximal fraction of active neurotransmitters can occur in each regime. In addition to that, assuming that the presynaptic neuron is spiking following a Poisson distribution, we showed that the equations for time average of the trajectory are the same as the map under periodic presynaptic stimulus, admitting the same equilibrium points. These results can contribute to understand how communication in the brain is mediated by synapses under regular or irregular stimulus.

The paper is organised as follows. In Section 2, we introduced the synaptic model considered in this work. In Section 3, we showed the analytical and numerical results. In the last section, we draw our conclusions.

## Methods

In short-term plasticity, the effective synaptic conductance changes in time depending on the neuron spike historic. The relevant parameters in synaptic dynamics are the presynaptic neuron firing frequency, the percentage of neurotransmitters released, and the time constants of the synapse. All these parameters in the model take into account the fact that there is a finite amount of neurotransmitters in each synapse and that they are not always available in the same quantity over time. Based on this, to study the synaptic dynamics, we considered the phenomenological model proposed in [22]. In this model, each directional synaptic connection from a presynaptic neuron [44] is represented by the set of ODEs

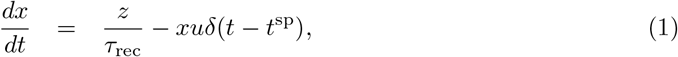

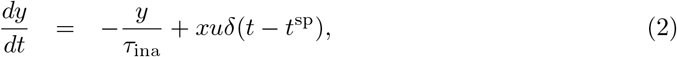

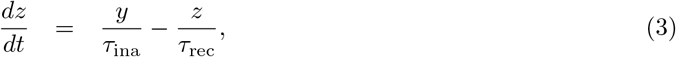

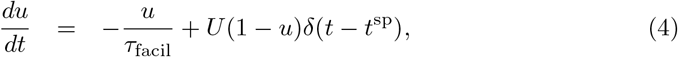

where *x, y*, and *z* represent the fractions of neurotransmitters in the recovered (available), active, and inactive state (refractory or recovering), respectively. *u* corresponds to the fraction of available resources (*x*) used in each presynaptic spike becoming active. In Eq (1), *τ*_rec_ is the recovery time for inactive neurotransmitters. In Eq (2), *τ*_ina_ is the decay time for neurotransmitters from active to inactive state. In Eqs (1) and (2), the Dirac delta function moves the fraction *xu* of neurotransmitter from the recovery state to the active one at the instant which a spike is considered in the model, namely *t*^sp^. We consider that when *t* = *t*^sp^, the delta Dirac function can be approximated by the Kronecker delta so that *δ*(*t* − *t*^sp^) = 1, otherwise *δ*(*t* − *t*^sp^) = 0. In Eq (3), the amount of recovering neurotransmitters depends on the inactivation and recovery process. In Eq (4), *τ*_fac_ is the time for the synapses to return from the facilitation regime. As *τ*_fac_ approximates to zero, less facilitation is exhibited in the model. *U* describes how *u* value increases and is associated with the percentage of available neurotransmitters which are released due to each spike. In this work, we fixed the parameters *τ*_rec_ = 800 ms, *τ*_ina_ = 3 ms, and *τ*_fac_ = 1000 ms [22]. Thus, we studied the parameters *U* and *t*^sp^, where *t*^sp^ is related to the mean spike frequency *F* and mean period *T* = 1*/F*. We consider the initial condition *x* = 1 and *y* = *z* = *u* = 0, which corresponds to a synapse with no recent activity and neurotransmitters totally recovered.

Fig 1 (a) shows a schematic representation of the neurotransmitters in the synapse by means of the variables *x, y*, and *z*. When a spike is considered in the model, the recovered neurotransmitters *x* can be released in the synaptic cleft, assuming an active state *y* that effectively will generate a current in the postsynaptic neuron. After the activation, these neurotransmitters are inactivated staying in the *z* variable until be in the recovered state *x* again and restart the cycle.

**Fig 1.**
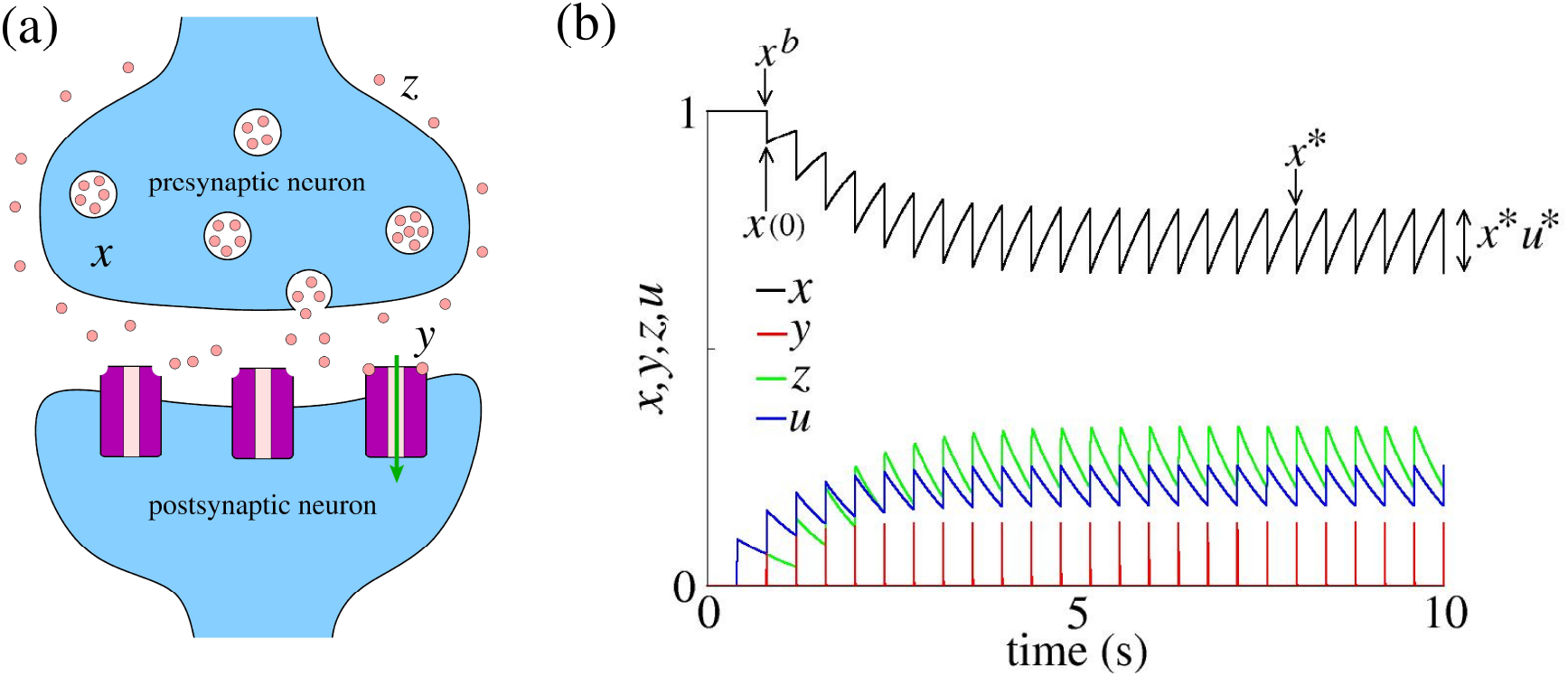
(a) Schematic representation of the neurotransmitter states in a synaptic connection. *x, y*, and *z* stand by recovered, active, and inactive neurotransmitters, respectively. (b) Variable evolution *x, y, z, u* overtime for the facilitation regime. *x*^*b*^ and *x*(0) stand by the *x* variable immediately before and after the spike event, respectively. *x*\* stands the fixed point of *x* after the transient time immediately before the spike event in the periodic spike regime. *x*\**u*\* represents the amount of neurotransmitters removed from the recovered state *x* and added to active state *y*. The same notation is used to identify the values of the variables before and after the spike event, as well as the fixed point after the transient. In Figure (b), we fixed *U* = 0.1 and *F* = 2.5 Hz.

## Results

### Analytical approximation, definitions and notations

To improve our understanding of the synaptic model, we search for an analytical approximation. We also introduce some notations and definitions. The variables *x, y*, and *u* are continuous in time when the evolution is considered between two sequential spikes events, from the time immediately after a spike event until the last time immediately before the next spike event. We assume *t*′ = *t* − *t*^sp^ = 0 to represent the time immediately after the spike of the neuron where *δ*(*t*′) = 1. Fig 1 (b) shows the time evolution of the variables, where *x*^*b*^ correspond to the value of *x* immediately before the spike (*t*^sp^). *x*(0) and *x*^*a*^ represent both the value of *x* immediately after a spike event (*t* = *t*^sp^), with the notation *x*(0) being used to handle the variables describing the evolution of the system of ODEs. Such notation is extended for other variables, *y, z*, and *u*. In this way, it is possible to relate the variable value before and after each spike to be considered in all variables by

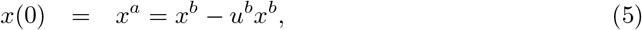

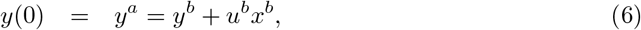

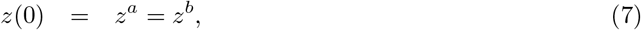

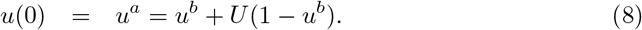

We seek a solution between two sequential spike events for the set of differential Eqs (1-4). We note that a general solution for *y*(*t*′) and *u*(*t*′) is independent of the other variables, resulting in

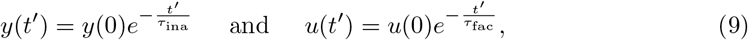

where *t*′ ∈ [0, *t*^sp^], *t*^sp^ being the time interval terminating just before the next spike happens. As can be observed, given a certain initial condition of these two variables, *y*(0) and *u*(0), the time evolution of *y* and *u* is determined until just before the next spike event. Since we have a solution for *y*(*t*′) and *z*(*t*′), it is possible to find an approximation for the solution of Eq (3) which is given by

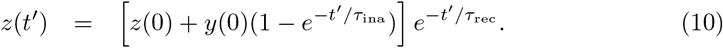

In the same way, as we have done for *z*(*t*′), we determine the solution of Eq (1) as

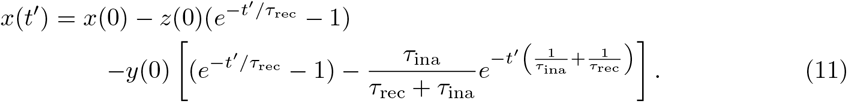

Taking into account Eqs (5-8) and the following definitions

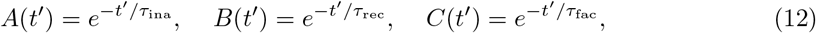

and

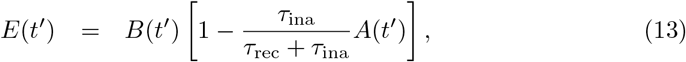

we can write Eqs (9), (10), and (11) as

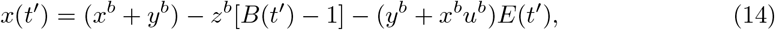

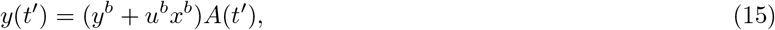

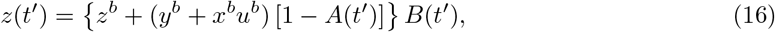

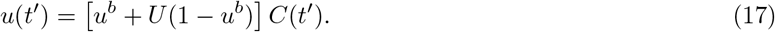

### Synaptic map model - Periodic regime

For periodic presynaptic spikes with a certain frequency *F*, the period between two spikes is given by *T* = 1*/F* and we set *t*′ = *t* − *t*^sp^ = *T*. The map is constructed by relating the value of the variables at the time *t*′ = *T* (immediately before the second spike)

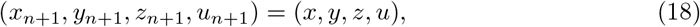

with those of the variables at the time *t*′ = 0 (immediately before the first spike)

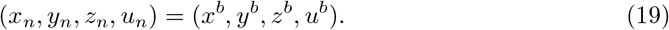

From Eqs (14-17), we obtain the map for the synapse

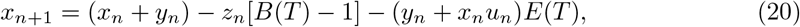

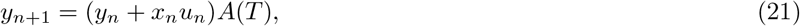

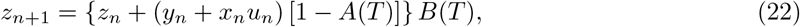

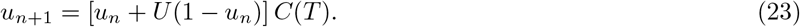

### Fixed point solutions

For periodic spikes, after a certain transient time, the equilibrium point represented by *x*\*, *y*\*, *z*\* and *u*\* is achieved. The equilibrium point is obtained by solving the system

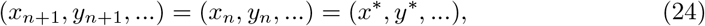

which lead us to

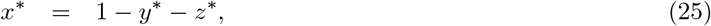

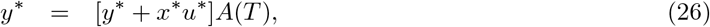

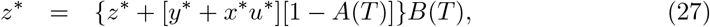

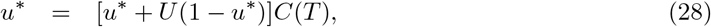

and finally

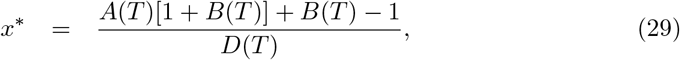

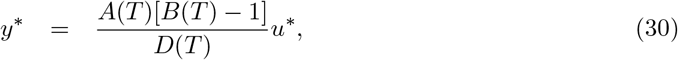

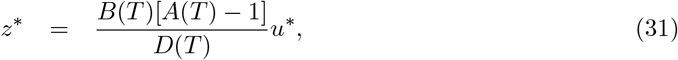

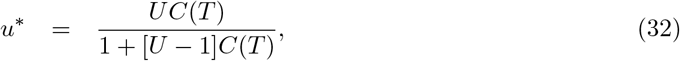

where

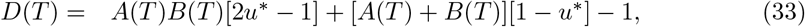

and *A, B* and *C* are defined in Eqs (12), for *t*′ = *T*.

### Approximation for low frequency

In the numerical simulation, we considered a small value of the time for inactivation given by *τ*_ina_ = 3 ms. Such value generates a fast decay in the *y* variable, so that we can approximate *y*_*n*_ to zero in Eqs (20), (21), and (22). In this case, we also can approximate *A*(*T*) to 0 and rewrite the map of Eqs (20-22) as

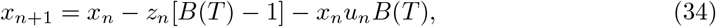

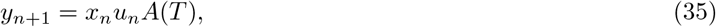

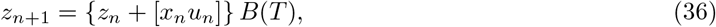

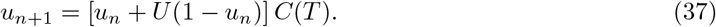

Since we can neglect *A*(*T*) for the inactivation time constant *τ*_ina_ = 3 ms and low frequencies, and noticing that *B*(*T*) and *C*(*T*) for the time constants *τ*_rec_ = 800 ms and *τ*_fac_ = 1000 ms have typically a far from zero value for low frequencies, we can rewrite the fix points as

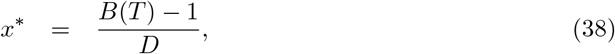

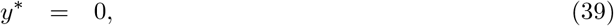

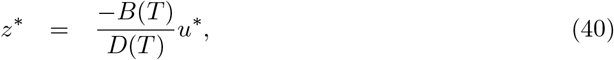

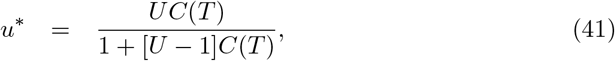

where

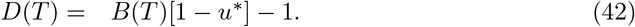

### Numerical analysis

By a numerical approach, we study the synaptic regimes as the function of the constant spike frequency *F* and the amount of neurotransmitter released associated with the parameter *U*. We demonstrate numerically the correctness of our analytical derivations for the map and its equilibrium points of active neurotransmitters.

To classify the synaptic regimes we considered the evolution of *y* due to the fact that it represents the effective fraction of neurotransmitters that is transmitted from the presynaptic to the postsynaptic neuron [22]. To mentioning, the synaptic current induced in the postsynaptic neuron is given by

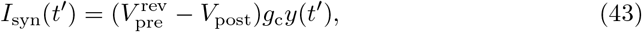

where 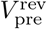 is the synaptic reversal potential associated with the presynaptic neuron type (excitatory or inhibitory), *V*_post_ is the potential of the postsynaptic neuron, *g*_c_ is the maximal synaptic conductance in the chemical synapse, and *y*(*t*′) is the fraction of active neurotransmitters released by the presynaptic neuron [7, 22, 45]. This quantity described by the model is the effective amount of neurotransmitters released due to each spike event, which in considered case has amplitude

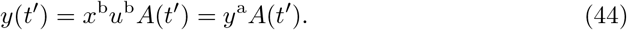

This value represents the fraction of neurotransmitters in the active state that will arrive in the receptors. Figs 2 (a-c) show examples of the three different regimes for periodic spikes, namely (a) facilitation, (b) biphasic, and (c) depression. The red line represents the fraction of active neurotransmitters, *y*(*t*), while the green points, *y*^a^, represent the maximal value of *y* due to each spike which describes the value of *y* immediately after the spike event. Based on this value, we define the three synaptic regimes previously mentioned. Facilitation corresponds to the synaptic regime where the value of active neurotransmitters only increases due to each spike or remains with an equal intensity after the transient period. Depression is the case where the value of active neurotransmitters only decrease due to each spike or remain at the same value after a transient time. Biphasic is associated with the synaptic regime where there is an increase and decrease in the value of active neurotransmitters. *y*_max_ represents the maximal value of *y* over time, while *y*_fin_ indicates the final amplitudes of *y* after the transient time. In Figs 2 (a-c), we considered *F* = 2.5 Hz, Fig (a) *U*=0.1 for facilitation, Fig (b) *U* =0.4 for biphasic, and Fig (c) *U*= 0.8 for depression regime. The mentioned parameters are indicated in Fig 2 (d) by white square, circle, and triangle, respectively.

**Fig 2.**
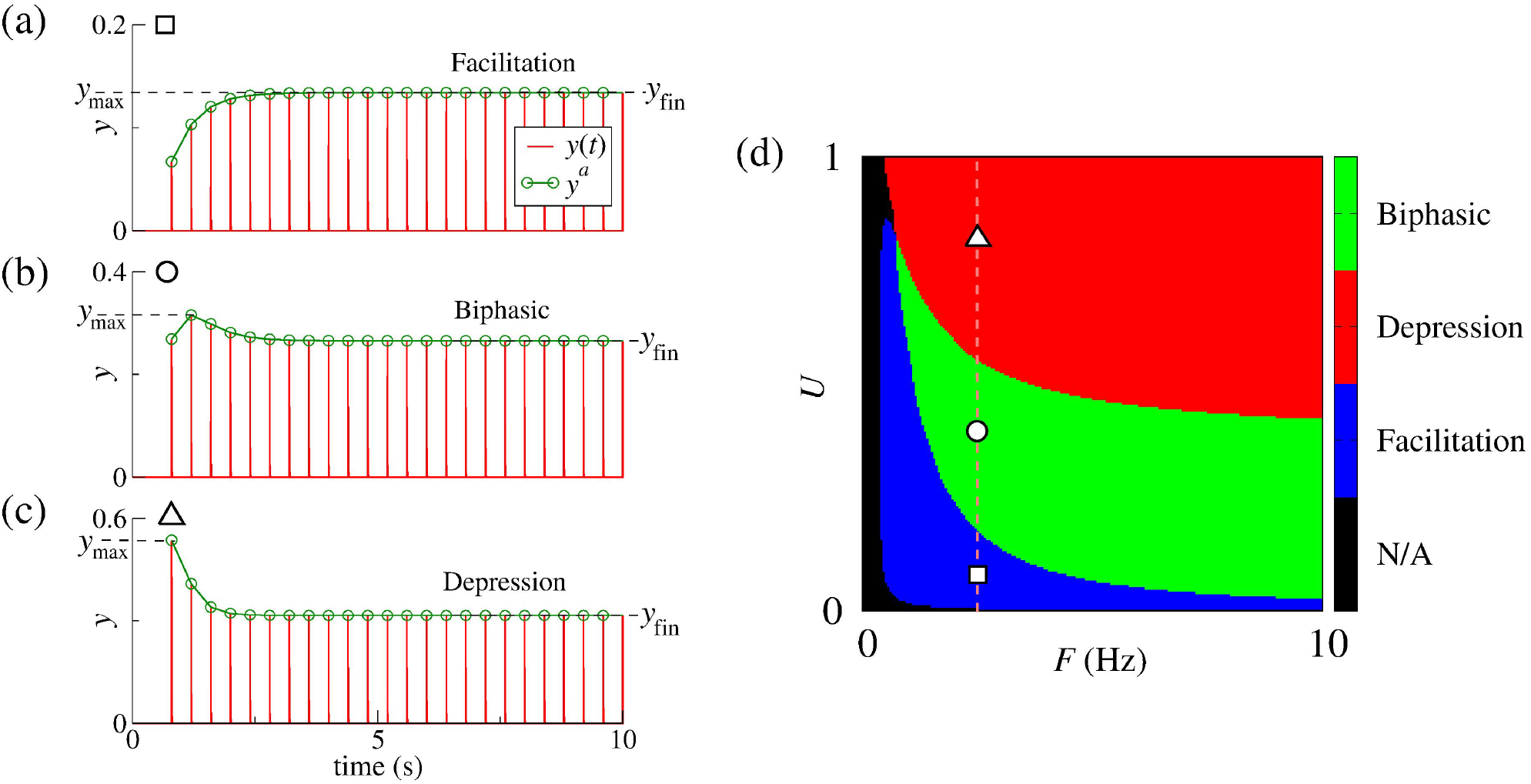
(a-c) Dynamics of the active neurotransmitters in the synapse for three different regimes, (a) facilitation, (b) biphasic, and (c) depression. Red lines show the dynamics of active neurotransmitters represented by the *y* variable, and the green line with circles represents the *y* amplitude immediately after the spike event, *y*^*a*^ = *y*(0). In Figs (a-c), we consider *F* =2.5 Hz, (a) *U* =0.1 for facilitation, (b) *U* = 0.4 for biphasic, and (c) *U* = 0.8 for depression regime. In the figures (a-c), the maximal value of *y* overtime is identified as *y*_max_ while the final amplitude of *y* is represented by *y*_fin_. (d) Regimes found in the dynamics synapses: facilitation, depression and biphasic. The facilitation and depression regimes are identified by means of Eqs (45) and (46). If both conditions of Eqs (45) and (46) are satisfied over time we identified the biphasic regime. The parameters *U* and *F* considered in Figs (a-c) are indicated in Fig (a).

Fig 2 (d) displays the synaptic regimes in the parameter space of the fraction of neurotransmitters released and the periodic spike frequency, *U* and *F*, respectively. As can be seen in the figure, the synaptic regimes are dependent on the parameters. The facilitation regime is found for the lowest frequency or smallest probabilities of neurotransmitter release, or both conditions. Depression is mainly found in the highest values of neurotransmitter release. Biphasic regime appears between the two previously cited regimes. In the map, we can identify the different regimes taken into account if some conditions are satisfied. In the facilitation regime, the amplitude of *y* variable will always increase or be equal to the final value *x*\**u*\* if

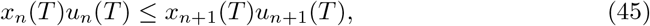

for *n >* 1 with fixed *U* and *T*, otherwise, for the depression regime, the amplitude of *y* variable will reduce or be equal to the final value *x*\**u*\*, if the condition

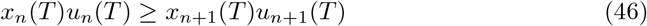

is satisfied for *n >* 1 with fixed *U* and *T*. If the Eqs 45 and 46 are satisfied in different iterations for *n >* 1, and fixed *T* and *U* values, the synapse exhibits a biphasic regime. If the fraction of neurotransmitter released or frequency assumes values very near to zero, the value of active neurotransmitters behaves as a linear dynamics where there is no change on this value (very small frequencies) or very close to zero (*U* close to zero), consequently, it is not possible to identify the regime in the parameter space. We identify such dynamics as “N/A” (not applicable) since the considered methodology does not identify such regimes, once conditions in Eqs (45) and (46) become close to equalities.

Fig 3 (a) shows the time evolution of the variables given by the model described by the ODE taking into account only the time immediately before the spike event. The figure also exhibits the value of *y* immediately after considering the spike event (*y*^*a*^) due to its importance in synaptic communication. The value of the fixed points of the map are determined by Eqs (38-41). The black dashed lines indicate the calculated fixed points *x*\*, *y*\*, *z*\*, and *u*\*, as well the final value of *y* immediately after the spike event, *y*_fin_. The value of *y*_fin_ is given by

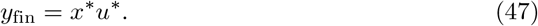

As it can be observed, the calculated fixed points agree with the evolution of the dynamics by Eqs (1-4), as well as the synaptic map model. Figs 3 (b) and (c) show the maximal and final value of *y*, where *y*_fin_ is obtained after the transient time. In Fig 3 (b), the maximal value corresponds to the highest value of *y* due to its entire time evolution given the initial condition of the synapse in the rest. The maximal values in the temporal series are found for the highest *U* and *F* values. However, these highest values of *y* might appear only briefly, cases observed in depression and biphasic regimes. For facilitation, the maximal *y* values correspond to *y*_fin_ and are exhibited after the transient time. As can be seen in Fig 3 (c), the final values of *y* do not appear for large frequencies, but there is a range of frequencies where *y*_fin_ is higher. For such a range of frequency, the higher values of *U* leads to higher *y*_fin_.

**Fig 3.**
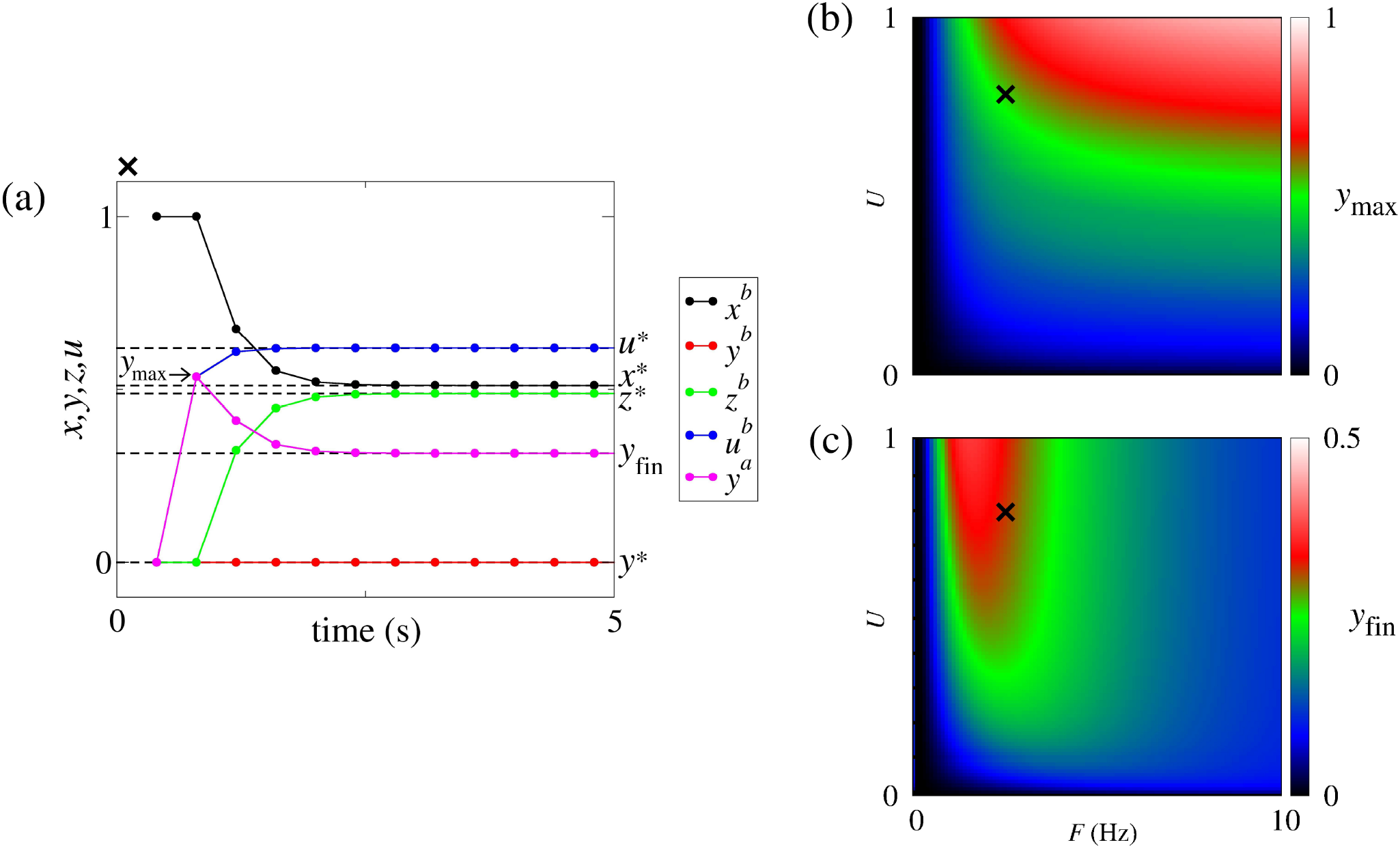
(a) Time evolution of the variable values immediately before considering the spike event on the synaptic model described by the EDO (*x*^*b*^, *y*^*b*^, *z*^*b*^, *u*^*b*^). As the *y* variable is responsible for generating the current in the postsynaptic neuron, the value of *y* immediately after considering the spike event (*y*^*a*^) and *y*_max_, are also indicated in the figure. The black dashed lines represent the calculated equilibrium points, *x*\*, *y*\*, *z*\*, and *u*\* of the discrete map, as well the final value *y*_fin_. The parameters *U* = 0.8 and *F* = 2.5 Hz are indicated in figures (b) and (c) by a black X. (b) Maximal *y* values in the parameter space *U* × *F*. *y*_max_ is obtained as the maximal value of *y* in the entire temporal evolution of the model. (c) *y*_fin_ in the equilibrium point, which is the *y* amplitude immediately after each spike past the transient time.

### Maximal fraction of active neurotransmitters

We study the maximal fraction of active neurotransmitters for a synapse initially on the rest. Figure 4 (a-d) shows the *y*^a^ dynamics for (a) depression, (b,c) biphasic, and (d) facilitation regime, identifying the *y*_max_ by a blue square. For depression, the maximal values occurs for the *n* = 1 while for biphasic regime occurs for *n* ≥ 2. Figure 4 (e) show the *n* value to find *y*_max_ in the synaptic map considering initial condition of the synapse on the rest state. The *n* value to obtain the *y*_max_ varies on the parameter space *F* × *U*.

**Fig 4.**
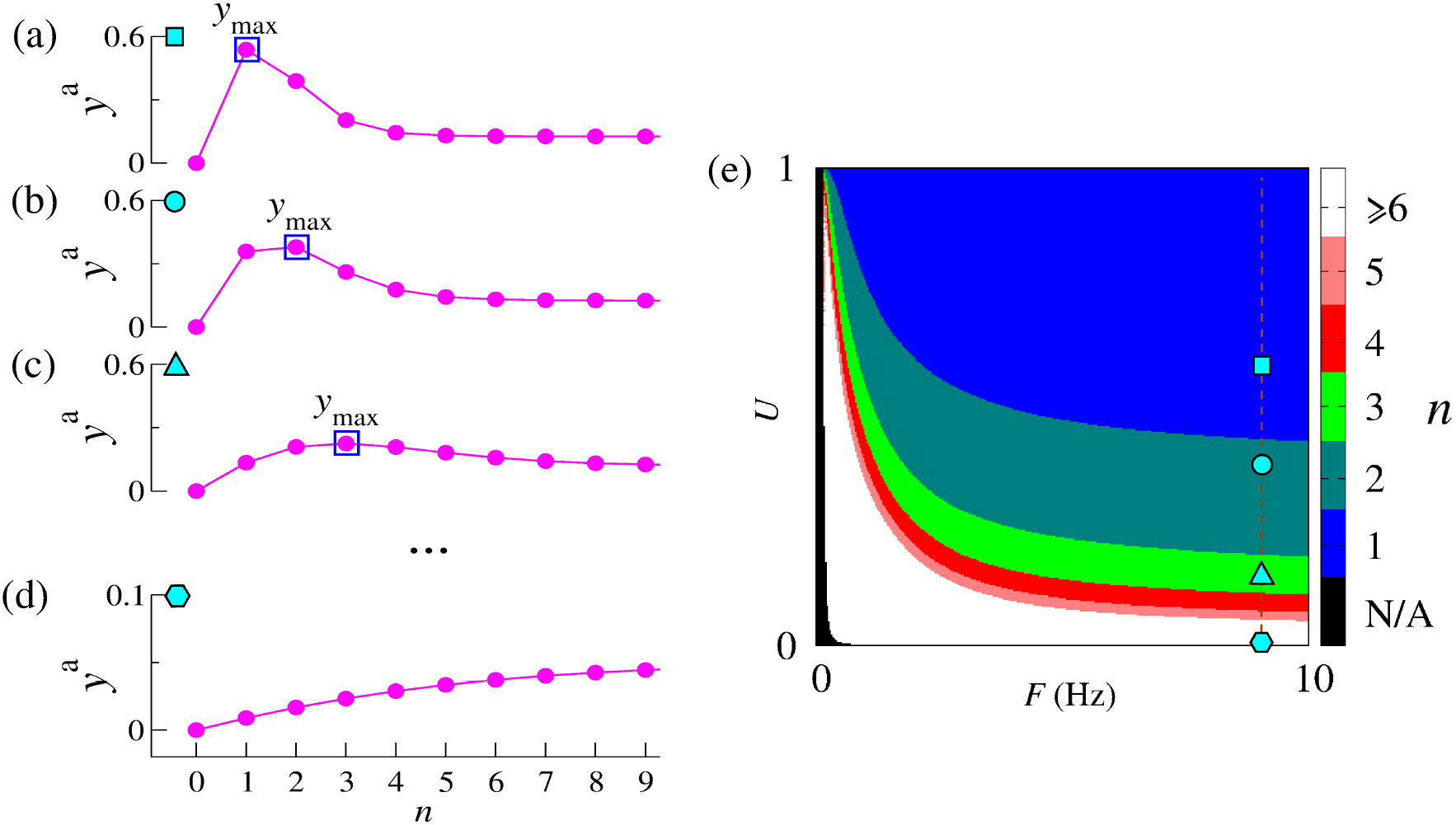
*y* variable after the spike (*y*^*a*^) for different values of the parameter of *U* identifying the maximal value (*y*_max_). In the Figure (a-d), we fix *F* =9 Hz, (a) *U* =0.6, *U* =0.4, (c) *U* = 0.15 and (d) *U* =0.01. Figure (e) shows the number of iterations to find the maximal value of *y* from the rest state of synapse.

To analytically determine the maximal values for the three synaptic regimes, we consider the following initial condition of the synapse on rest,

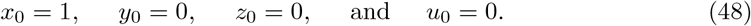

These values represent the state variables immediately before the first spike. One map iteration takes them to the states immediately before the second spike, given by

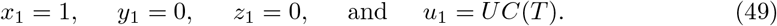

For the depression regime showed in Figure 4 (a), the maximal value of *y* is given by Eq. (15) for *t*′=0, and therefore equal to

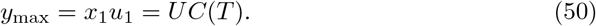

For the biphasic regime showed in Figure 4 (b,c), the maximal *y* value is also provided by Eq. (15) for *t*′=0, however depending on the parameters that maximal is only reached after a certain number of spikes. So,

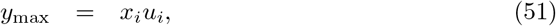

where *i* representing the number of spikes applied to the initial condition at rest can assume values equal to 2, 3, 4, …. Abusing the notation and dropping the argument (*T*) of *A*(*T*), *B*(*T*), and *C*(*T*), for simplicity, the variable states immediately before the second spike is given by

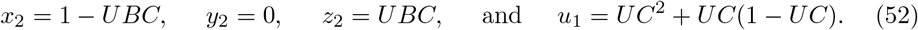

Inputting these values into Eq. (15) provides the potential maximal value after 2 spikes of the neuron (*i* = 2) given by

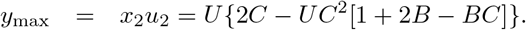

This maximal value for *i* = 2 is predominant in the parameter space *F* × *U* for the biphasic regime. The larger the value of *i*, the smaller the area in the parameter space, showing that it is less likely to be observed. Notice that to calculate the maximal values analytically, the larger the value of *i* the larger the degree of the polynomial associated with the solution seek. So, we only calculate maximal values up to *i*=2.

Finally, for the facilitation regime showed in Figure 4 (d), the maximal value of *y* is asymptotically increasing towards the equilibrium point and can be calculated by using the values from the equilibrium points in Eqs. (38) and (41) into Eqs. (14) and (17), and then plugging these values into Eq. (15) for *t*′ = 0.

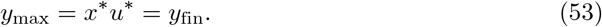

### Synaptic map model - Poissonian regime

In this section, we consider non-periodic spike times given by a homogeneous Poissonian process to generate the time of spikes *t*^sp^. Different from the periodic case, the time interval *T* between two spikes is not constant but rather is described by a Poisson process and thus assumes different values for each iteration. Namely, for *N* + 1 iterations, we have *T*_0_, *T*_1_, *T*_2_, …, *T*_*N*_, where *N* is considered as a large integer value. Just showing the iteration for these time intervals for the *y* variable described by Eq (21), we have

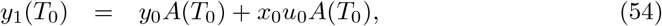

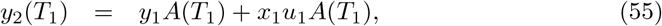

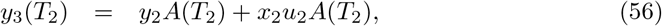

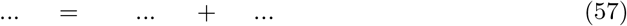

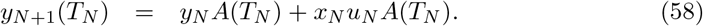

Considering the sum of all the terms in the equations and dividing by *N* + 1 to obtain a temporal average, we find

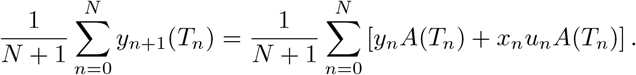

Identifying the mean values by

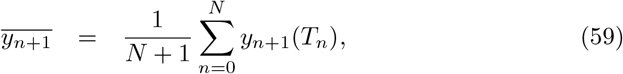

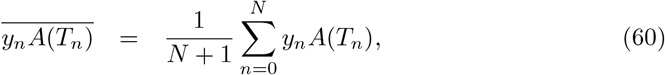

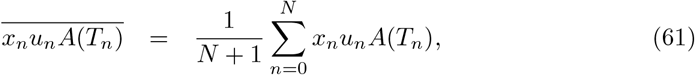

we can write the expression of the temporal average for the *y* variable as

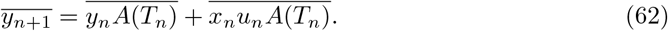

Taking into account that the product of the mean is equal to the mean of products, we can rewrite the last expression as

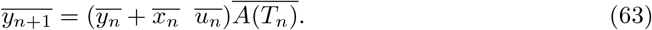

To determine the value of 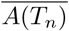, we consider the power series expansion for exponential function in the case that *T*_*n*_ *<< τ*_fac_, obtaining the approximation

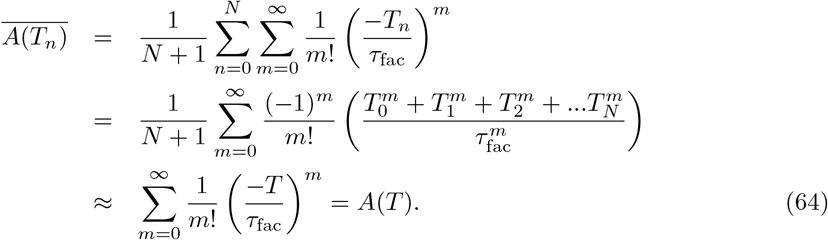

Using the same calculations for the other map variables in Eqs (20), (22), and (23), for a homogeneous Poisson process, we obtain the map evolution of the temporal average given by

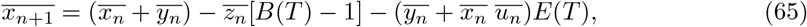

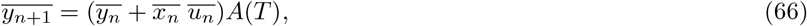

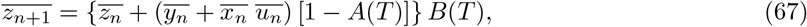

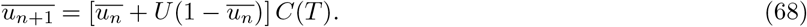

The temporal averaged map for the Poissonian presynaptic spikes exhibits the same evolution as the synaptic map for presynaptic neurons spiking periodically, since the equations are the same. These equations for the temporal averaged map admit a solution for the equilibrium point if

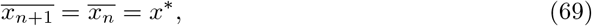

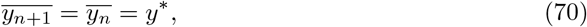

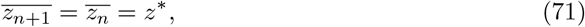

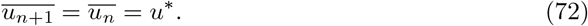

Since Eqs (20-23) are the same as Eqs (65-68), the equilibrium points of the periodic stimulus is the same for the Poissonian stimulus. Furthermore, the time average value of *xu* immediately after spikes for both the Poissonian and periodic spikes tends to product *x*\**u*\*. Our result suggests that the average behaviour of a synapse forced by Poissonian spikes is determined by the dynamics of synapses driven by periodic spikes.

## Conclusion

In this paper, we study short-term plasticity considering the model proposed by Tsodyks et al. [22]. We focus our analysis on the synaptic regimes facilitation, depression, and biphasic that emerge as a function of the frequency and fraction of released neurotransmitters. Depression regime is mainly dependent on the percentage of released neurotransmitters, while facilitation is observed for low frequencies or small amounts of neurotransmitter release, or for both cases. The biphasic regime is found between the facilitation and depression, being a combination of both regimes.

Our main result was to obtain an approximated solution for the set of differential equations and derive a map where the synaptic dynamics can be understood in terms of the time intervals between the spike events. From this map, we determined analytically the equilibrium points for periodic spiking regimes, allowing for the determination of the asymptotic values of active neurotransmitters as a function of the frequency, the probability release of neurotransmitters, and the time constants. We also determined the expected maximal and asymptotic values for the three synaptic regimes. We observe that the highest value of active neurotransmitters occur by a brief time period from the rest regime in the depression and biphasic regime, while for facilitation regime, the maximal values are the asymptotic ones.

Furthermore, we obtain a temporal average map when the time intervals between spikes follows a Poisson distribution and show that the equilibrium point for such a configuration is the same found for the periodic spikes. This result suggests that the time average dynamics of a stochastic driven synapse emulating presynaptic spikes by a complex neural network is regulated by the periodically driven synapses.

## Acknowledgments

This study was possible by partial financial support from the following Brazilian government agencies: FAPESP (2020/04624-2, 2022/05153-9, 2022/13761-9).

## Author Contributions

**Conceptualization:** Paulo R. Protachevicz, Antonio M. Batista, Iberê L. Caldas, and Murilo S. Baptista.

**Methodology:** Paulo R. Protachevicz and Murilo S. Baptista.

**Formal analysis:** Paulo R. Protachevicz and Murilo S. Baptista.

**Investigation:** Paulo R. Protachevicz and Murilo S. Baptista.

**Validation:** Paulo R. Protachevicz.

**Formal analysis:** Paulo R. Protachevicz and Murilo S. Baptista.

**Funding acquisition:** Iberê L. Caldas.

**Writing – original draft:** Paulo R. Protachevicz and Murilo S. Baptista.

**Writing – review & editing:** Paulo R. Protachevicz, Antonio M. Batista, Iberê L. Caldas, and Murilo S. Baptista.

## References

1. Di Maio V, Santillo S, Ventriglia F. Synaptic dendritic activity modulates the single synaptic event. Cogn Neurodyn. 2021; 15(2): 279–297.

2. Pereda AE. Electrical synapses and their functional interactions with chemical synapses. Nat Rev Neurosci. 2014; 15: 250–263.

3. Connors BW, Long MA. Electrical synapses in the mammalian brain. Annu Rev Neurosci. 2004; 27(1): 393–418.

4. Caire MJ, Reddy V, Varacallo M. Physiology, Synapse. In: StatPearls. Treasure Island (FL): StatPearls Publishing; 2022.

5. Bear MF, Connors BW, Paradiso MA, Michael A. Neuroscience: Exploring the Brain. Jones and Bartlett Publisher, Inc. Fourth Edition; 2020.

6. Suszkiw JB. Synaptic transmission. In: Sperelakis N, editor. Cell Physiology Source Book, fourth ed., Elsevier; 2012. pp. 563–578.

7. Gerstner W, Kistler WM, Naud R, Paninski L. Neuronal dynamics: From single neurons to networks and models of cognition and beyond. Cambridge University Press; 2014.

8. Roth A., van Rossum Mcw. Modeling Synapses, in Erik De Schutter (ed). Computational Modeling Methods for Neuroscientists. Cambridge, MA. MIT Press Scholarship Online; 2009.

9. Sterrat D, Graham B, Gillies A, Willshaw D. The synapses. Principles of computational modelling in neuroscience. Cambridge University Press; 2012.

10. Tauffer L, Kumar A. Short-term synaptic plasticity makes neurons sensitive to the distribution of presynaptic population firing rates. eNeuro. 2021; 8(2): 0297.

11. Roberts PD. Synaptic dynamics: Overview. In: Jaeger, D., Jung, R. (eds). Encylopedia of Computational Neuroscience. Springer, New York;2014

12. Citri A, Malenka RC. Synaptic plasticity: Multiple forms, functions, and mechanisms. Neuropsychopharmacology. 2008; 33: 18–41.

13. Deperrois N, Graupner M. Short-term depression and long-term plasticity together tune sensitive range of synaptic plasticity. PLoS Comput Biol. 2020: 16(9): e1008265.

14. Bliss TVP, Cooke SF. Long-term potentiation and long-term depression: a clinical perspective. Clinics. 2011; 66:3–17.

15. Kemp A, Manahan-Vaughan D. Hippocampal long-term depression and long-term potentiation encode different aspects of novelty acquisition. PNAS. 2004; 101(21): 8192–8197.

16. Howell RD, Pugh JD. Biphasic modulation of parallel fibre synaptic transmission by co-activation of presynaptic GABAA and GABAB receptors in mice. J Physiol. 2016; 1;594(13): 3651–66.

17. Cho S, Li G-L, von Gersdorff H. Recovery from short-term depression and facilitation is ultrafast and Ca+2 dependent at auditory hair cell synapses. J Neurosci Res. 2011; 31(15), 5582–5692.

18. MacLeod KM, Horiuchi TK, Carr CE. A role for short-term synaptic facilitation and depression in the processing of intensity information in the auditory brain stem. J Neurophysiol. 2007; 97: 2863–2874.

19. Buonomano D V Decoding temporal information: A model based on short-term synaptic plasticity. J Neurosci Res. 2000; 20(3): 1129–1141.

20. Deng P-Y, Klyachko VA. The diverse function of short-term plasticity components in synaptic computation. Commun Integr Biol. 2011; 2(5): 543–548.

21. Abbott LF, Regehr WG. Synaptic computation. Nature. 2004; 431: 796–803.

22. Tsodyks M, Uziel A, Markram H. Synchrony generation in recurrent networks with frequency-dependent synapses. J Neurosci Res. 2000; 20: 1–5.

23. Rotman Z, Deng P-Y, Klyachko VA. Short-term plasticity optimizes synaptic information transmission. J Neurosci Res. 2011; 31(41): 148000–14809.

24. Citri A, Malenka RC. Synaptic Plasticity: Multiple forms, functions, and mechanisms. Neuropsychopharmacol Rep. 2008; 33: 18–41.

25. Cortes JM, Desroches M, Rodrigues S, Veltz R, Munñoz MA, Sejnowski TJ. Short-term synaptic plasticity in the deterministic Tsodyks-Markram model leads to unprecdictable networks dynamics. PNAS. 2013; 110(41): 16610–16615.

26. Markram H, Wang Y, Tsodyks M. Differential signaling via the same axon of neocortical pyramidal neurons. PNAS. 1998; 95(9): 5323–5328.

27. Fortune ES, Rose GJ. Short-term synaptic plasticity as a temporal filter. Trends Neurosci. 2001; 24(7): 381–385.

28. Barroso-Flores J, Herrera-Valdez MA, Galarraga E, Bargas J. Models of short-term synaptic plasticity. Adv Exp Med Biol. 2017; 1015: 41–57. Springer International Publishing AG.

29. Blitz DM, Foster KA, Regehr WG. Short-term synaptic plasticity: A comparison of two synapses. Nature Publishing Group. 2004; 5: 630–640.

30. Jackman SL, Regehr WG. The mechanisms and functions of synaptic facilitation. Neuron. 2017; 94(3): 447–464.

31. Barak O, Tsodyks M. Persistent activity in neural networks with dynamic synapses. PLOS Comput Biol. 2007; 3(2): e35.

32. Mondal Y, Pena RFO, Rotshein HG. Temporal filters in response to presynaptic spike trains: interplay of cellular, synaptic and short-term plasticity time scales. J Comput Neurosci. 2022; 50: 395–429.

33. Kass RE, Ventura V. A spike-train probability model. Neural Comput. 2001; 13(8): 1713–20.

34. Deger M, Cardanobile S, Helias M, Rotter S. The poisson process with dead time captures important statistical features of neural activities. BMC Neurosci. 2009; 10: P110.

35. Simen P, Balci F, deSouza L, Cohen JD, Holmes P. A model of interval timing by neural integration. J Neurosci Res. 2011; 31(25): 9238–9253.

36. Burkitt AN. A review of the integrate-and-fire neuron model: I. Homogeneous synaptic input. Biol Cybern. 2006; 95: 1–19.

37. Burkitt AN. A review of the integrate-and-fire neuron model: II. Inhomogeneous synaptic input and network properties. Biol Cybern. 2006; 95:97–112.

38. Cinlar E. Introduction to stochastic processes. Courier Corporation; 2013.

39. Gabbiani F, Cox SJ. Mathematics for Neuroscientist. Academic Press, Elsevier; 2010.

40. Ladenbauer J, McKenzie S, English DF, Hagens O, Ostojic S. Inferring and validating mechanistic model of neural microcircuits based on spike-train data. Nat Commun. 2019; 10: 4933.

41. Pena RFO, Zaks MA, Roque AC. Dynamics of spontaneous activity in random networks with multiple neuron subtypes and synaptic noise. J Comput Neurosci. 2018;45: 1–28.

42. Droste F, Lindner B. Up-down-like background spiking can enhance neural information transmission. eNeuro. 2017; 4(6): 0282–17.

43. Mazzoni A, Broccard FD, Garcia-Perez E, Bonifazi P, Ruaro ME, Torre V. On the dynamics of the spontaneous activity in neuronal networks. PLos One. 2007; 2(5): e439.

44. Bertolotti E, Burioni R, di Volo M, Vezzani A. Synchronization and long-time memory in neural networks with inhibitory hubs and synaptic plasticity. Phys Rev E. 2017; 95: 012308.

45. Protachevicz PR, Borges FS, Lameu EL, Ji P, Iarosz KC, Kihara AH, Caldas IL, Szezech Jr, J., Baptista MS, Macau EEN, Antonopoulos CG, Batista AM, Kurths J. Bistable Firing Pattern in a Neural Network Model Front Comput Neurosci. 2020; 13(19).

